# Cell cycle regulators control mesoderm specification in human pluripotent stem cells

**DOI:** 10.1101/632307

**Authors:** Loukia Yiangou, Rodrigo A. Grandy, Sanjay Sinha, Ludovic Vallier

## Abstract

Mesoderm is one of the three germ layers produced during gastrulation from which muscle, bones, kidneys and the cardiovascular system originate. Understanding the mechanisms controlling mesoderm specification could be essential for a diversity of applications, including the development of regenerative medicine therapies against diseases affecting these tissues. Here, we use human pluripotent stem cells (hPSCs) to investigate the role of cell cycle in mesoderm formation. For that, proteins controlling G1 and G2/M cell cycle phases were inhibited during differentiation of hPSCs into lateral plate, cardiac and presomitic mesoderm using small molecules or by conditional knock down. These loss of function experiments revealed that CDKs and pRb phosphorylation are necessary for efficient mesoderm formation in a context-dependent manner. Further investigations showed that inhibition of the G2/M regulator CDK1 decreases BMP signaling activity specifically during lateral plate mesoderm formation while reducing FGF/ERK1/2 activity in all mesoderm subtypes. Taken together, our findings reveal that cell cycle regulators direct mesoderm formation by controlling the activity of key developmental pathways.

---

Gastrulation represents an essential stage of early development when the three primary germ layers endoderm, mesoderm and ectoderm are formed. Of particular interest for the current study, mesoderm specification occurs through a mesendoderm progenitor which is shared with endoderm. Mesoderm is then patterned in different subpopulations depending of their anteroposterior location in the primitive streak where a gradient of Nodal and BMP signaling and interplay with WNT and FGF leads to the formation of different cell types (1–3). At the very posterior end of the primitive streak, high BMP4 and low Nodal pattern extraembryonic mesoderm and the blood lineage, followed by anterior lateral plate mesoderm (1, 4, 5). The lateral plate mesoderm lineage gives rise to cardiovascular cell types such as smooth muscle cells and endothelial cells. At the more anterior part moderate levels of Nodal and BMP4 pattern cardiac mesoderm, which gives rise to cardiomyocytes, the main cell type constituting the heart (4, 6). In the middle anterior primitive streak, paraxial (presomitic) mesoderm is formed (7) which gives rise to bone, cartilage and skeletal muscle and requires WNT signaling. Understanding the mechanisms directing the specification of these different types of mesoderm could have a broad implication for developmental biology but also in the context of diseases affecting stem cell differentiation. Nonetheless, studying these mechanisms at the molecular level remains challenging *in vivo* due to technical and ethical limitations in human.

Human pluripotent stem cells (hPSCs) provide a powerful alternative since they can proliferate almost indefinitely while maintaining the capacity to differentiate efficiently into the three germ layers (8). Thus, hPSCs have been used to uncover mechanisms directing germ layer specification (9–11). Of particular interest, studies have shown key functions for the cell cycle machinery in the specification of endoderm versus ectoderm and exist from the pluripotent state. Indeed, G1 and G2/M transition regulators have been shown to play a key role in pluripotency maintenance and cell fate decisions of hPSCs by controlling transcription factors, signaling pathways and epigenetic regulators (12–16). More precisely, knockdown of CDK2 results in cell cycle arrest, decreased expression of pluripotency markers and differentiation towards extraembryonic lineages (17). Similarly, abrogation of cyclin D1/2/3 results in loss of pluripotency and differentiation towards the mesendoderm lineage (13) indicating a direct role of cyclins and CDKs in the maintenance of pluripotency and cell identity. Furthermore, siRNA mediated knockdown of CDK1 results in changes in cell morphology, decrease in pluripotency marker expression, accumulation of DNA damage and mitotic deficiencies (18).

At the epigenetic level, histone modification H3K4me3 has been shown to be more abundant on developmental genes in the G1 phase of the cell cycle. Interestingly, the histone methyltransferase catalysing this modification called MLL2, was also shown to be higher in the late G1 phase and enriched on promoters of the cell cycle regulated genes *SOX17* and *GATA6*. The recruitment of MLL2 on these developmental genes was shown to be mediated by phosphorylation of MLL2 by CDK2, establishing a role of cell cycle regulators in regulating epigenetic processes in hESCs (15).

Moreover, the cell cycle machinery and specifically the cyclin D-CDK4/6 complex has a vital role in guiding endoderm formation through regulation of the Nodal/Activin signaling pathway effector Smad2/3. Cyclin D also acts as a transcriptional regulator independently of its role in the cell cycle and signaling regulation. ChIP-sequencing analyses have shown that cyclin D binds to and recruits transcriptional corepressor and coactivator complexes onto developmental gene loci, thus regulating their transcription and ultimately cell fate decisions (14).

Most of these studies, if not all, have been performed in the context of pluripotency while the role of the cell cycle in guiding differentiation especially mesoderm specification is elusive. Here, we decided to address this question by taking advantage of protocols allowing differentiation of hPSCs into different mesoderm subtypes including lateral plate mesoderm (LPM), cardiac mesoderm (CM) and presomitic mesoderm (PSM) (19, 20). The corresponding culture system was used to investigate the role of G1 and G2/M cell cycle regulators in guiding mesoderm specification, especially the role of the cell cycle machinery in regulation of key developmental signaling pathways such as BMP, WNT and FGF. Inhibition of both G1 and G2/M cell cycle regulators by small molecules blocked mesoderm subtype formation with different efficacy and in a context-dependent manner. Additional analyses into the molecular mechanisms responsible for this phenotype, revealed that inhibition of the G2/M regulator CDK1 decreased BMP activity during LPM formation while reducing FGF/ERK activity in all mesoderm subtypes studied, thereby blocking differentiation. Our results demonstrate that cell cycle regulators are essential for the early stage of mesoderm formation and that this function is achieved through regulation of key developmental signalling pathways such as FGF/ERK. This knowledge will help to improve protocols for generating mesoderm cells *in vitro* and could also be relevant for the development of new therapies promoting tissue regeneration.

## RESULTS

### Characterisation of mesoderm subtypes generated from hPSCs

In this study, we took advantage of established protocols for differentiating hPSCs into different mesoderm subtypes. Specifically, we took advantage of chemically defined culture conditions to drive differentiation of hPSCs into cardiac (CM), lateral plate (LPM) and presomitic (PM) mesoderm. These methods rely on growth factors known to direct mesoderm specification *in vivo* (20–22). As a result, hPSCs differentiation follows a natural path of development including the production of cells closely resembling cells arising along the antero-posterior axis of the primitive streak during development. In sum, hPSCs were induced to generate LPM, CM and PSM mesoderm for 36 hours followed by the addition of another cocktail of growth factors and small molecules to generate functional cell types such as smooth muscle cells, cardiomyocytes and chondrocytes (Fig. 1*A*) (20, 22, 23). During induction of all mesoderm subtypes, we observed a decrease in pluripotency marker expression such as *NANOG* and upregulation of pan-mesoderm marker *BRACHYURY* (or *T*) (Fig. 1*B-G*). LPM induction was confirmed by the increase in *NKX2.5* expression at day 5 (Fig. 1*B, C*) and further differentiation toward smooth muscle cells was validated by the expression of calponin (*CNN1*) and transgelin (*TAGLN*) at day 17 (Fig. 1*B, C*). During CM induction we observed expression of *EOMES* at day 1.5. CM identity was confirmed by the high expression of *NKX2.5* at day 6 while further differentiation resulting in beating cardiomyocytes expressed the genes *ACTN1* (coding for the microfilament protein α-Actinin) and *TNNT2* (coding for cardiac Troponin T) (Fig. 1*D, E*). Finally, PSM induction was associated with CDX2 expression at day 1.5 followed by chondrocyte differentiation as shown by the expression of the cartilage matrix proteins collagen 2a (or *COL2A1*) and aggrecan *(ACAN*) (Fig. 1*F, G*). Immunostaining analyses showed PAX3 expression in PSM while alcian blue staining confirmed production of proteoglycans such as aggrecan by terminally differentiated chondrocytes (Fig. 1*G*). Taken together, these results reinforce previous results by showing the robustness of our protocols to drive differentiation of hPSCs into different types of mesodermal progenitors.

**FIGURE 1.**
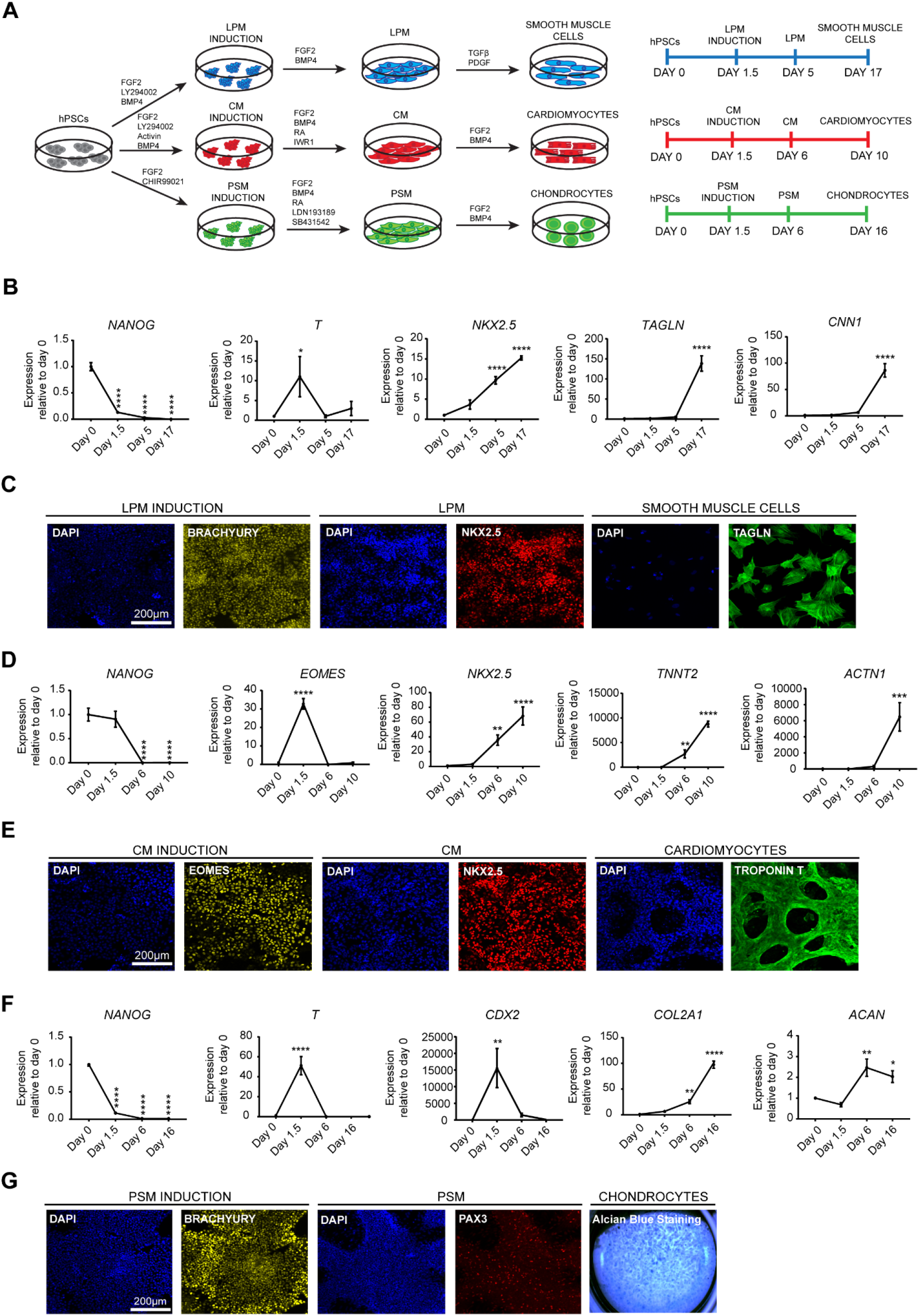
Production of mesoderm subtypes from hPSCs. *A*, Schematic overview of the differentiation protocol. LPM was induced by FGF2, the PI3K inhibitor LY294002 and BMP4 and then SMCs were generated using TGFβ and PDGF-BB. CM was induced by Activin, FGF2, the PI3K inhibitor LY29400 and BMP4 and cardiac cells were obtained by inhibiting WNT signaling in the presence of BMP4, FGF2 and Retinoic Acid (RA). PSM was induced by FGF2 and the WNT signalling agonist CHIR99021. PSM was then generated using FGF2, RA and dual inhibition of TGFβ and BMP4 signalling using the small molecules SB431542 and LDN193189 respectively. Chondrocyte differentiation was induced by FGF2 and BMP4. *B*, RT-qPCR analysis for expression of pluripotency (*NANOG)* and mesoderm markers (*T*, *NKX2.5*, *TAGLN* and *CNN1*) during SMC differentiation. *C*, Immunostaining analysis for the expression of early mesoderm markers BRACHYURY, NKX2.5 and SMC marker TAGLN. Scale bar: 200µm. *D*, RT-qPCR analysis for the expression of pluripotency (*NANOG*) and mesoderm markers (*EOMES*, *NKX2.5*, *TNNT2* and *ACTN1*) during cardiomyocyte differentiation. *E*, Immunostaining analysis for the anterior primitive streak marker EOMES, CM marker NKX2.5 and cardiomyocyte marker Troponin T. Scale bar: 200µm. *F*, RT-qPCR analysis for expression of pluripotency (*NANOG*) and mesoderm (*T*, *CDX2*, *COL2A1* and *ACAN*) markers during chondrogenic differentiation. *G*, Immunostaining analysis for the expression of early mesoderm marker BRACHYURY, PSM marker PAX3 and Alcian blue staining of chondrocytes differentiation. Scale bar: 200µm. Error bars represent ±SEM (n=6). Ordinary one-way ANOVA test followed by Dunnett’s test for multiple comparisons was performed. *p<0.05, **p<0.01, ***p<0.001, ****p<0.0001.

### Inhibition of G1 and G2/M cell cycle regulators blocks induction of mesoderm subtypes in a context-dependent manner

To explore the importance of cycle machinery in mesoderm specification, we next investigated the effect of the inhibition of G1 and G2/M regulators on differentiation. For that, we used small molecule inhibitors for CDK4/6 (PD-0332991), CDK2 (roscovitine), phosphorylation of retinoblastoma protein (RRD-251) and CDK1 (RO-3306 (Fig. 2*A*). Of note, these small molecules are commonly used to study the function of cell cycle regulators in a diversity of systems. hESCs were induced to differentiate into the three mesoderm subtypes LPM, CM and PSM in the presence of the cell cycle regulators inhibitors for 36 hours and the resulting cells were harvested for gene expression and immunocytochemistry analyses (Fig. 2*B*).

**FIGURE 2.**
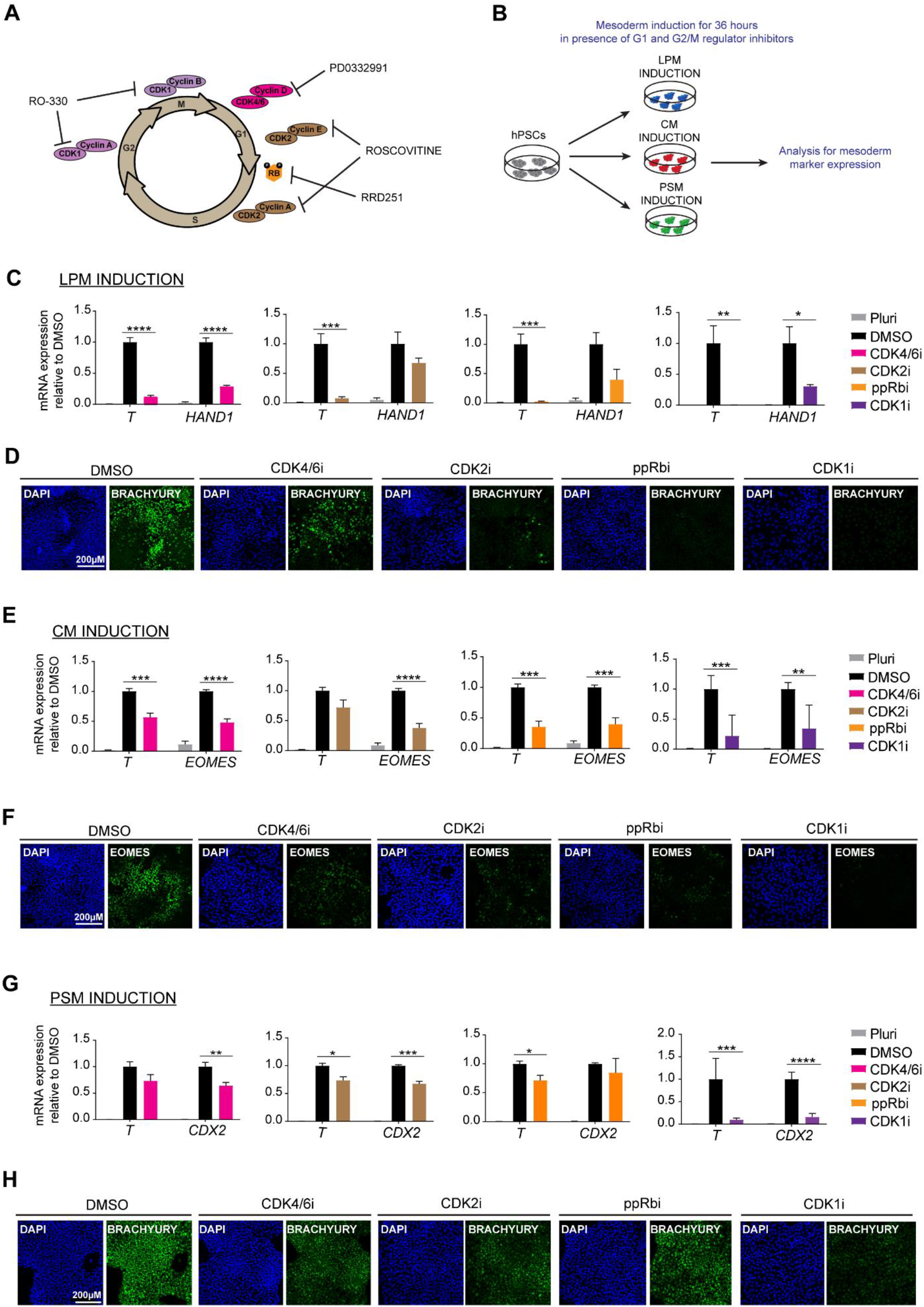
Inhibition of cell cycle regulators blocks induction of mesoderm subtypes. *A*, Schematic showing the action of the different cell cycle regulator inhibitors used in the study. *B*, Schematic overview of experimental setup to investigate the role of cell cycle regulators in mesoderm specification. For each experiment hESCs were differentiated in the absence (DMSO) or presence of inhibitor. *C*, RT-qPCR analysis for the expression of mesoderm markers (*T* and *HAND1*) during LPM induction (n=3). *D*, Immunostaining analysis for BRACHYURY expression during LPM induction. Scale bar: 200µm. *E*, RT-qPCR analysis for the expression of mesoderm markers (*T* and *EOMES*) during CM induction (n=5). *F*, Immunostaining analysis for EOMES expression during CM induction. Scale bar: 200µm. *G*, RT-qPCR analysis for the expression of mesoderm markers (*T* and *CDX2*) during PSM induction (n=5). *H*, Immunostaining analysis for the expression of BRACHYURY during PSM induction. Scale bar: 200µm. Error bars represent ±SEM. Unpaired t-test was performed. *p<0.05, **p< 0.01, ***p<0.001, ****p<0.0001.

Concerning LPM differentiation, CDK2, ppRb and CDK1 inhibition were associated with strong downregulation of *T* expression (Fig. 2*C*). Moreover, the posterior primitive streak marker *HAND1* was also downregulated upon inhibition of all regulators, with the strongest effect being observed during inhibition of CDK4/6 and CDK1 (Fig. 2*C*). The inefficient formation of mesoderm was also validated at the protein level where a reduction in expression of BRACHYURY was observed upon CDK4/6 inhibition whereas inhibition of CDK2, ppRb and CDK1 resulted to complete loss of BRACHYURY expression (Fig. 2*D*). Of note, downregulation of the pluripotency markers *OCT4* and *NANOG* was not stopped suggesting that inhibition of cell cycle regulators did not block differentiation of hPSCs (Fig. S1*A*).

Similarly, CM specification was strongly altered upon inhibition of all cell cycle regulators, as shown by the reduction in *T* and *EOMES* expression (Fig. 2*E*, *F*). Interestingly, inhibition of CDK2, pRb and CDK1 resulted in maintenance of *OCT4* and *NANOG* expression during the differentiation process (Fig. S1*B*). Thus cell cycle regulators may be necessary to exit pluripotency during differentiation of specific types of mesoderm.

PSM induction was also inefficient upon inhibition of the cell cycle regulators. Expression of *T* was more significantly reduced upon inhibition of CDK2, ppRb and CDK1 and expression of *CDX2* upon inhibition of CDK4/6, CDK2 and CDK1 (Fig. 2*G*). Immunostaining analysis for BRACHYURY expression confirmed RT-qPCR results (Fig. 2*H*). Expression of pluripotency genes did not significantly differ between DMSO and inhibitor-treated cells with the exception of CDK1 inhibition which led to the maintenance of *OCT4* and *NANOG* expression during the differentiation process (Fig. S1*C*). This suggests that CDK1 could be necessary for the downregulation of these genes during PSM formation.

Taken together, these results show that cell cycle regulators are essential for mesoderm formation and subtypes specification. Of note, the strong effect seen on *T* expression suggests that these regulators may be involved during the early stage of mesoderm specification corresponding to primitive streak formation. Furthermore, CDKs and the pRb protein could also be necessary for the exit from pluripotency as suggested by the pattern of expression for *OCT4* and *NANOG*.

### Inhibition of G1phase regulators does not affect the activity of BMP, WNT and FGF signaling pathways

Based on our previous results showing that CDK4/6 could control the activity of Activin/Nodal signaling (13), we hypothesised that regulators of G1 phase including CDK4/6, CDK2 and pRb could control signaling pathways such as BMP, WNT, and FGF which are known to direct mesoderm specification. To confirm this possibility, we studied the effect of G1 CDK inhibitors on the activity of these signaling pathways during the specification of different types of mesoderm. Effect on BMP signaling was analysed in the context of LPM and CM formation since BMP4 is used drive formation of these mesoderm subtypes while effect on WNT signaling was analysed during PSM formation since CHIR99021, a WNT agonist, is driving this mesoderm formation. Finally, effect on FGF signaling was primarily analysed in pluripotent cells to avoid interference with other pathways. Indeed, FGF2 is important for all three mesoderm subtypes LPM, CM and PSM and thus it was more informative to test the corresponding inhibitors in pluripotency conditions before further investigations during differentiation. Each mesoderm subtype was induced in the presence of the inhibitors for 36 hours with the exception of FGF signaling, for which hESCs were grown for 12 hours in the presence of inhibitors. Prior to harvesting for Western blot analysis, cells were freshly fed with media supplemented with the inhibitors. and subsequently harvested for Western blot analyses (Fig. 3*A*).

**FIGURE 3.**
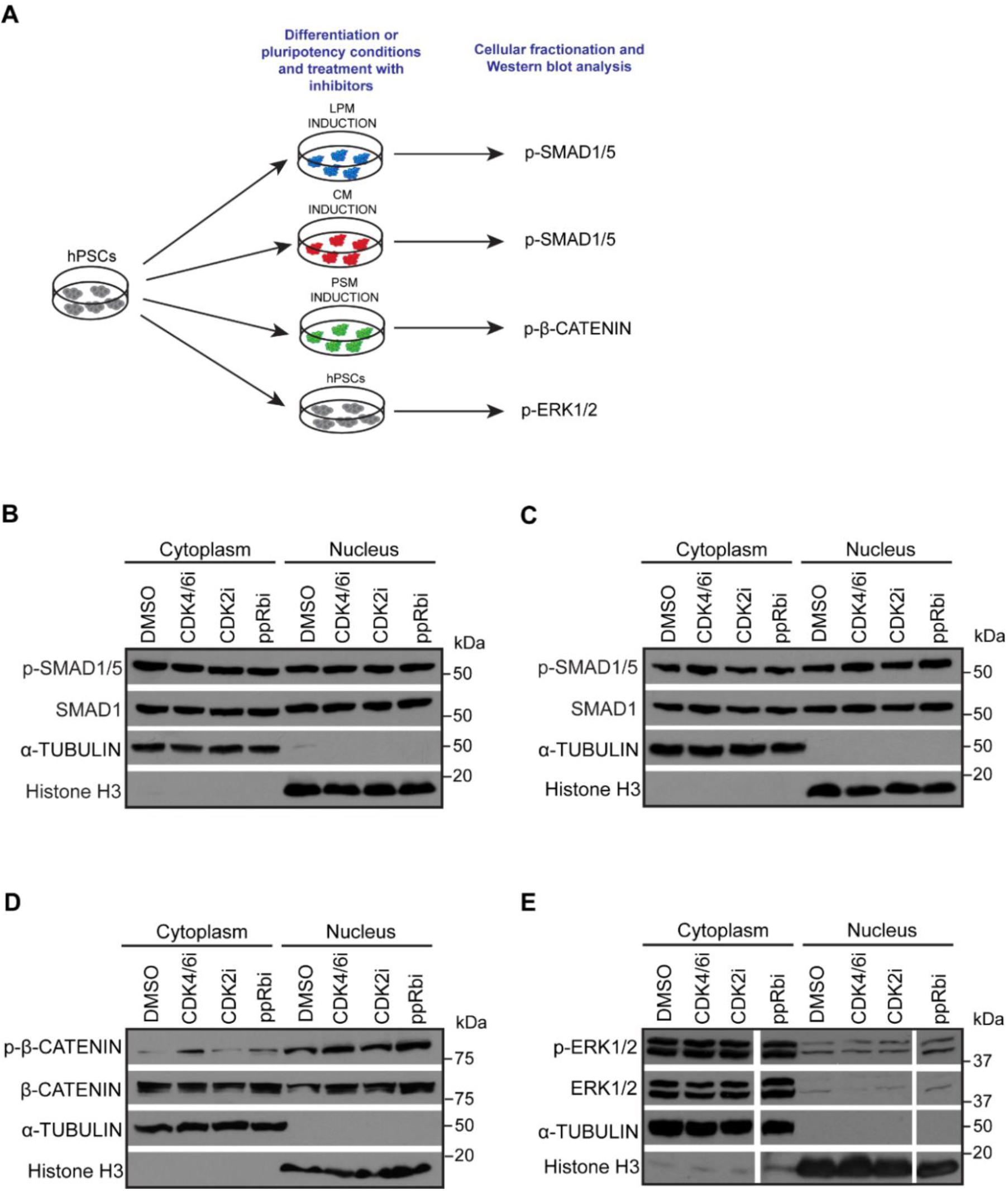
BMP, WNT and FGF signaling pathways are not affected by inhibition of G1 regulators. *A*, schematic overview of experimental setup to determine activity of BMP, WNT and FGF signaling upon treatment with G1 phase regulator inhibitors. *B-C*, Western blot analyses for phospho-SMAD1/5 and total SMAD1 to determine activity of BMP signaling in DMSO-treated vs cells treated with G1 regulator inhibitors during LPM (*B*) and CM induction (*C*). *D*, Western blot analyses for phospho-β-catenin and total β-catenin to determine activity of WNT signaling. *E*, Western blot analyses for phospho-ERK1/2 and total ERK1/2 to determine activity of FGF signaling. Blot represents samples run on the same gel. A-tubulin and Histone H3 were used as loading controls for the cytoplasm and the nuclear fractions respectively.

Despite several analyses and different conditions of treatment, inhibition of cell cycle regulators in these different culture conditions did not affect the levels of phospho-SMAD1/5 (Fig. 3B-C) neither the levels of phospho-β-catenin (Fig. 3*D*) or the level of phospho-ERK1/2 (Fig. 3*E*). Thus, G1 phase regulators do not appear to regulate BMP, WNT or FGF signaling, suggesting that the regulation of mesoderm formation could happen through alternative mechanisms.

### Inhibition of CDK1 decreases the activity of BMP and FGF signaling pathways

We next investigated the effect of CDK1 inhibition on the same signaling pathways using similar conditions. Interestingly, CDK1 inhibitor strongly reduced levels of phospho-SMAD1/5 (Fig. 4*A*) specifically during LPM differentiation but not induction of CM (Fig. 4*B*). Thus, CDK1 could control BMP signaling only in certain mesoderm subtypes. Further characterisation showed that WNT signaling pathway was not affected by CDK1 (Fig. 4*C*) while FGF/ERK1/2 signaling pathway was severely down regulated upon inhibition of CDK1 in pluripotency conditions (Fig. 4*D*) and even more strongly in all mesoderm subtypes. The most severe phenotype was observed in LPM and CM induction where loss of phospho-ERK1/2 was almost complete (Fig. 4, *E* and *F*) while this reduction was less severe in PSM (Fig. 4*G*). To confirm these results, we decided to genetically validate this phenotype by knocking-down CDK1 in hESCs. For that, we took advantage of the single-step optimized inducible knockdown (sOPTiKD) platform as previously described (24). The resulting CDK1 iKD-hESCs were treated with tetracycline for 5 days to induce the knockdown of CDK1 and subsequently differentiated into the three mesoderm subtypes in the presence of tetracycline. Western blot analysis confirmed the efficient knockdown of CDK1 with 80%, 60% and 90% decrease in expression in LPM, CM and PSM respectively (Fig. 4*H*, first panel). Western Blot analyses showed that decrease in CDK1 expression was associated with reduced ERK1/2 phosphorylation (Fig. 4*H*, second panel). The strongest effect was observed in LPM and CM induction whereas reduction in PSM was less severe, recapitulating the phenotype obtained by inhibiting CDK1 with small molecule. Taken together, these results demonstrate that CDK1 is necessary for the activity of BMP and FGF signaling during mesoderm specification.

**FIGURE 4.**
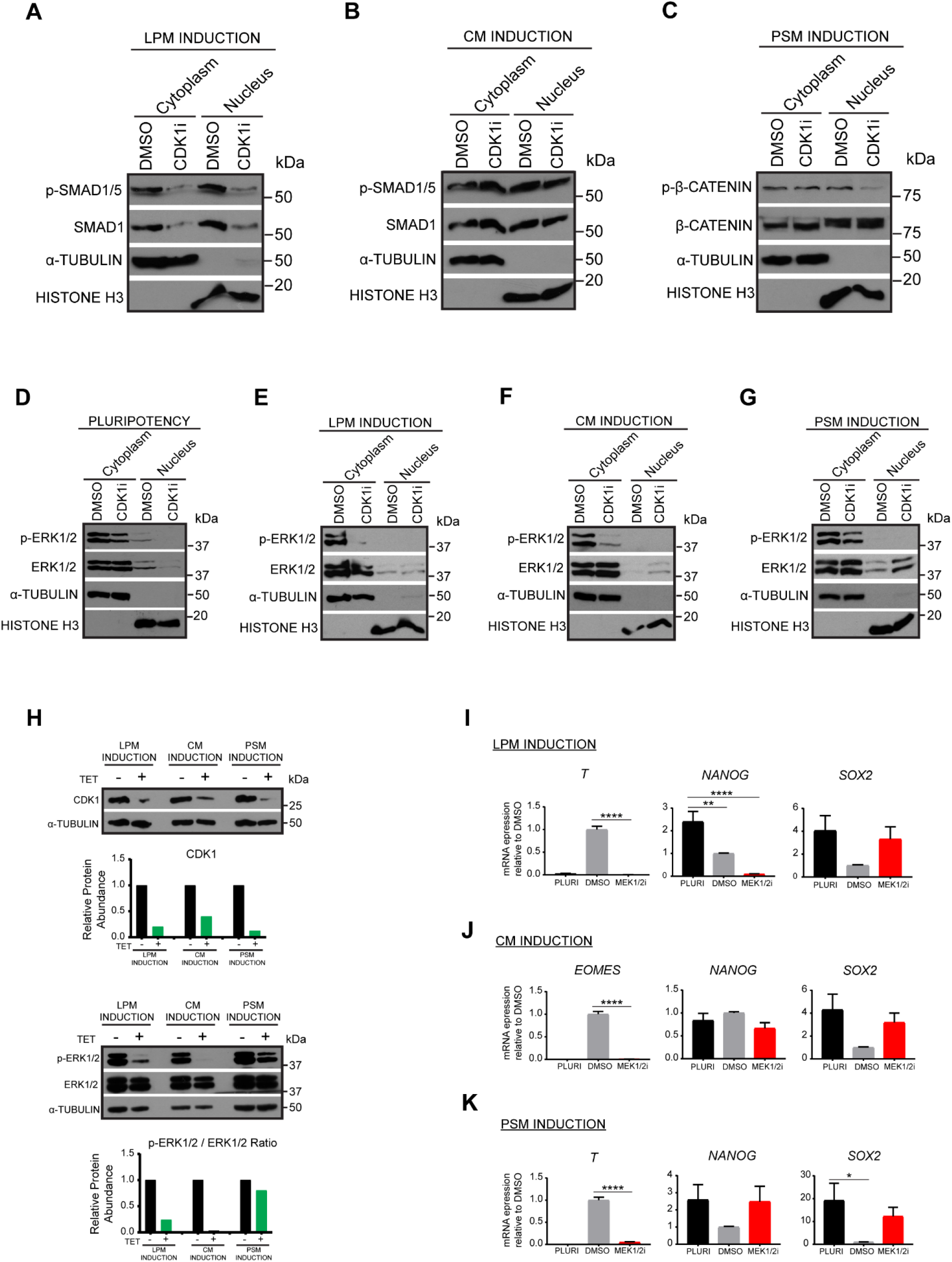
Inhibition of CDK1 decreases BMP and FGF signaling pathway activity. *A-B*, Western blot analyses for phospho-SMAD1/5 and total SMAD1 to determine activity of BMP signaling in DMSO-treated vs cells treated with the CDK1 inhibitor RO-3306 during LPM (*A*) and CM induction (*B*). *C*, Western blot analyses for phospho-β-catenin and total β-catenin to determine activity of WNT signaling. *D-G*, Western blot analyses for phospho-ERK1/2 and total ERK1/2 to determine activity of FGF signaling in pluripotency conditions (*D*) and during LPM (*E*), CM (*F*) and PSM induction (*G*). Α-tubulin and Histone H3 were used as loading controls for the cytoplasm and the nuclear fractions respectively. *H*, Western blot analysis of CDK1 iKD cells. Pluripotent cells were treated with tetracycline for 4 days prior to induction of differentiation. Following 36 hours of mesoderm subtype differentiation, cells were harvested for analysis of phospho-ERK1/2 and CDK1 expression. Graphs show densitometric quantitation of protein relative to loading control α-tubulin and normalised to –TET treatments. *I-K*, RT-qPCR analysis for expression of differentiation and pluripotency markers during induction of LPM (*I*), CM (*J*) and PSM (*K*). Error bars represent ±SEM(n=3). Ordinary one-way ANOVA test followed by Tukey’s test for multiple comparisons was performed. *p<0.05, **p< 0.01, ***p<0.001, ****p<0.0001.

### Inhibition of FGF/ERK1/2 signaling blocks induction of all mesoderm subtypes

our results also imply that ERK1/2 should be essential for mesoderm differentiation. To validate this hypothesis, we induced mesoderm subtype formation for 36 hours in the presence of the ERK/MEK1/2 inhibitor PD0325901. Strikingly, presence of this small molecule resulted in the complete loss of expression of *T* during LPM (Fig. 4*I*) and PSM induction (Fig. 4*K*) and loss of *EOMES* during CM induction (Fig. 4*J*). In addition, *SOX2* expression was up regulated in all the mesoderm types (Fig. 4*I-K*) while *NANOG* down regulation was only affected in PSM thereby recapitulating the results obtained with CDK1 inhibition. Taken together, these results demonstrate that mesoderm subtype induction is controlled by the interplays between CDK1 and FGF/ERK1/2 which are necessary for the induction of key mesoderm markers but also the down regulation of pluripotency factors.

## DISCUSSION

In this study, we investigated the role of G1 and G2/M cell cycle regulators in the specification of mesoderm subtypes. We used pharmacological inhibition of CDK4/6, CDK2 and CDK1 using the small inhibitors PD-0332991, roscovitine and RO-3306 respectively. Additionally, retinoblastoma protein (pRb) was maintained in its active state by inhibiting its phosphorylation with the small molecule RRD-251. Inhibition each cell cycle regulator disrupted specific mesoderm differentiation. CDK4/6 appears to be necessary for all the mesoderm subtypes thereby suggesting a general role for this regulator in inducing mesoderm formation. Interestingly, it was previously shown that CDK4/6 inhibits endoderm formation by blocking the nuclear import of SMAD2/3 (13). Thus, cell cycle regulators could be germ layer specific and context-dependent. In our mesoderm differentiation model, only anterior primitive-like and subsequent cardiac mesoderm is induced in the presence of Activin and high activity of SMAD2/3 is likely to perturb mesoderm induction. Thus, CDK4/6 could be necessary to safe guard mesoderm induction against ectopic Activin signaling activity.

Interestingly, CDK2 and phosphorylation of pRb also appear to be necessary for mesoderm specification. CDK2 is known to control phosphorylation of pRb during cell cycle progression and thus both regulators could interact during differentiation (Reviewed in Harbour and Dean, 2000). Accordingly, the most important effect of their inhibition is the loss of *T* expression specifically during LPM induction. These results also suggest that CDK2 and/or pRb could control the early stage of mesoderm induction corresponding to the primitive streak formation *in vivo*. Despite their functional importance, we were not able to establish a link between G1 phase regulators with signaling pathways controlling differentiation. Thus, CDK2/pRb could control alternative mechanisms necessary for mesoderm patterning. It has been previously proposed that cell cycle regulators and specifically CDK2 regulates epigenetic modifiers such as the MLL methyltransferase and guides it to developmental genes in the late G1 phase (15). This observation could also apply in our system and specifically it would be of interest to identify whether MLL2 is phosphorylated by CDK2 or CDK4/6 during mesoderm differentiation and whether there is recruitment of MLL2 to genes guiding mesoderm specification such as BRACHYURY whose expression has been shown to rely on-MLL2 (15).

In contrast to G1 cell cycle regulators, we observed a strong reduction in FGF/ERK1/2 signaling upon CDK1 inhibition in all mesoderm subtypes, and a reduction in BMP4/SMAD1/5 signaling in during LPM induction. This observation is consistent with the interplay of FGF and BMP4 signaling in hESCs which is known to be key for mesoderm differentiation (26, 27). Thus, CDK1 could be a link between the two signaling pathways, whose coordinated function is necessary for the expression of key mesoderm inducers such as *T*. FGF has been shown to regulate *T* expression in xenopus, zebrafish, chick and mouse embryos (28–33) and a correlation of FGF signaling and *T* expression was reported in human cancer cell lines (34). Thus, our results support *in vivo* and *in vitro* findings and suggest that a similar mechanism may be conserved in human stem cells.

Beyond differentiation markers, we also observed that inhibition of CDK2, ppRb and CDK1 could affect pluripotency markers. Decrease in *OCT4* and *NANOG* expression was accentuated during LPM induction and partially inhibited during CM induction. Thus, for some mesoderm subtypes, cell cycle regulators could fine-tune exit from pluripotency mechanisms in agreement with previous reports (12), while for other mesoderm subtypes they could regulate the expression of pluripotency factors, guiding differentiation (20, 35–37).

Our results, suggest an interesting link between pluripotency factor expression and upregulation of differentiation genes. It has been shown that during BMP-induced differentiation, NANOG expression is prolonged (through ERK signaling) and NANOG knockdown causes loss of *T* expression in hESCs (26). Additionally, functional studies showed that loss of NANOG reduces expression of basal levels of primitive streak genes and loss of OCT4 results in *T* decrease in hESCs. Conversely, overexpression of NANOG was shown to increase levels of primitive streak genes (35). Moreover, a recent study has identified that ERK2 and CDK1 both phosphorylate NANOG (38). This is an intriguing correlation suggesting that the two kinases could stabilise NANOG through phosphorylation in the G2/M phase in a cooperative manner. Considered together, these reports reinforce the data presented in the current study.

Importantly, validations of our finding in vivo remain challenging. Indeed, knockout of G1 cell cycle regulators results in viable embryos (39–43) while only the simultaneous knockout of CDK2 and CDK4 is embryonic lethal (44). These results suggest that G1 CDKs are not essential for embryo development at the early stages, possibly due to functional redundancy often witnessed among these regulators. On the other hand, knockout of CDK1 is essential for cell proliferation during early development and its absence causes embryonic lethality at E10.5 (45). CDK1 thus is considered a master regulator of cell cycle transition. Interestingly, the essential role of CDK1 has been further corroborated as it was shown to compensate for loss of CDK2, CDK4 and CDK6 in the cells by binding to cyclins E, A, B and D (46, 47), highlighting its essential role throughout development.

Of note, inhibition of CDK1 in our system during the differentiation process could be blocking cell cycle progression, thus contributing to the inefficient differentiation of the cells. While this being a possibility, studies from our lab have shown that blocking or slowing down cell cycle during differentiation using small molecule such as nocodazole does not prevent the induction of specific markers such as Brachyury (Grandy RA., et al., unpublished observation).

Importantly, we cannot exclude that CDK1 could control additional molecular mechanisms directing differentiation. Indeed, CDK1 is also known to control epigenetic modifiers such as the polycomb group protein enhancer of zeste homologue 2 (EZH2). This protein is a histone methyltransferase and catalyses H3K27me3 leading to gene silencing and has been shown to have a role in maintenance of pluripotency in PSCs, amongst its multiple other roles (48–50). Intriguingly, CDK1 was shown to control EZH2 activity through phosphorylation (51) while EZH2 inhibition was shown to be important for mesoderm formation in hESCs (52). Considered together, these previous studies suggest that CDK1 could coordinate epigenetic modification with signaling pathway activity during progression of mesoderm differentiation.

Taken together, our results show that cell cycle regulators are essential for mesoderm differentiation and exit from pluripotency. The mode of action of these regulators appears to be specific for each mesoderm subtype depending on the regulation of *T* by pluripotency factors and FGF/ERK1/2 signaling (Fig. 5). These mechanisms represent an additional step to understand the precise interplays between cell cycle machinery and differentiation.

**FIGURE 5.**
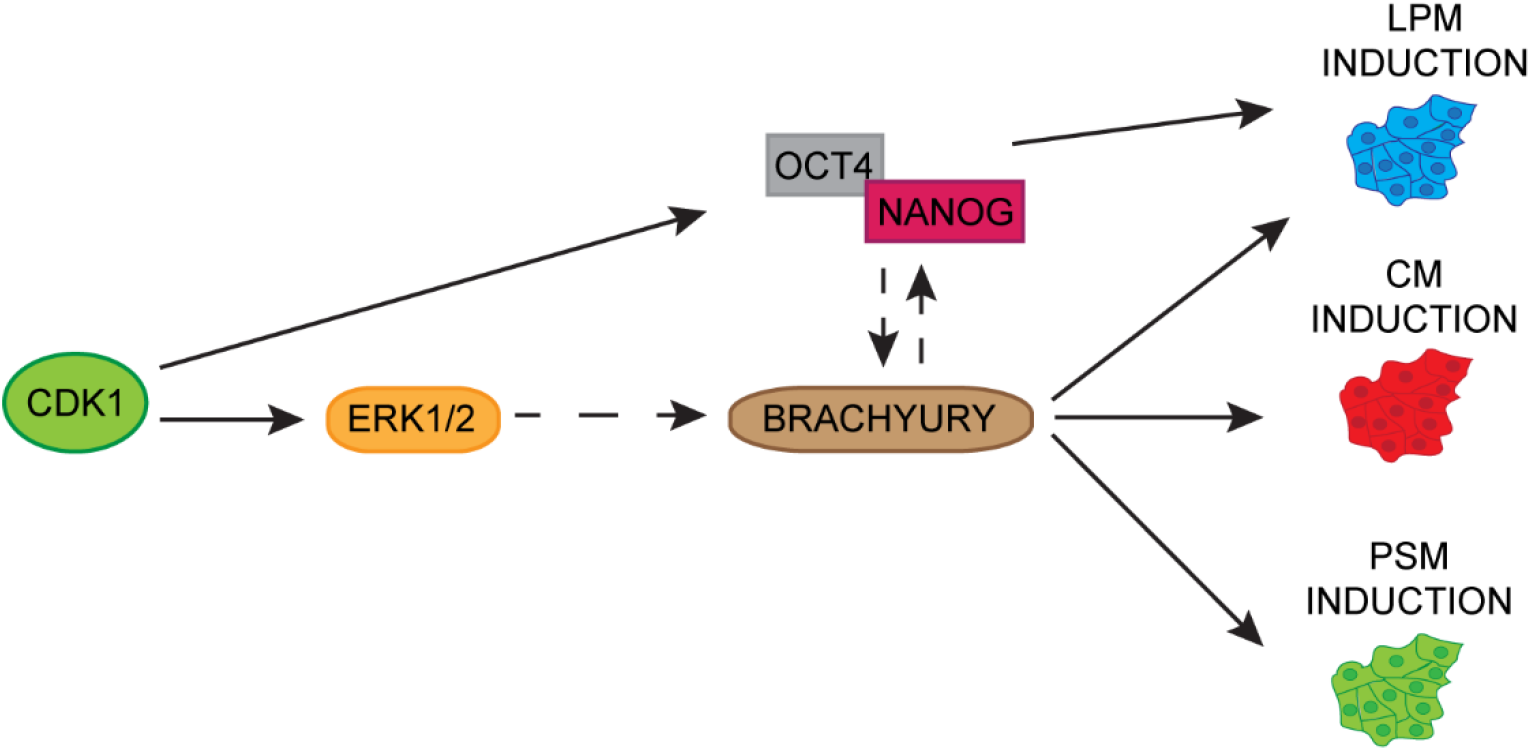
Model depicting the role of CDK1 in mesoderm specification. Schematic of main findings on role of CDK1 in mesoderm patterning. CDK1 regulates FGF/ERK1/2 signaling to control *BRACHYURY* expression and subsequent mesoderm differentiation. Additionally, CDK1 could be directly or indirectly regulating expression of OCT4/NANOG to drive LPM formation.

## EXPERIMENTAL PROCEDURES

### hESC culture and differentiation

H9 hESCs (WiCell, Madison, WI, USA) were cultured on vitronectin coated plates (10µg/ml, Stem Cell Technologies) in E6 media supplemented with 2ng/ml TGF-β (R&D) and 25ng/ml FGF2 (Dr. Marko Hyvönen, Cambridge University). Cells were maintained by weekly passaging using 0.5mM EDTA (Thermo Fisher Scientific). CDK1 iKD H9 hESCs cells were grown in the same conditions, supplemented with 1µg/ml puromycin for selection of antibiotic resistant cells. CDK1 knockdown was induced by adding 2µg/ml tetracycline hydrochloride (TET) (Sigma-Aldrich) to the culture medium 4 days prior to the start of differentiation.

The cells were differentiated into the three germ layers and functional cell types as previously described (20, 23). Mesoderm subtypes were generated in a 2-step protocol. For lateral plate mesoderm (LPM) formation, cells were cultured for 36 hours in CDM-PVA supplemented with 20ng/ml FGF2, 10µM LY294002 (Promega) and 10ng/ml BMP4 (R&D). Subsequently cells were cultured for 3 days in CDM-PVA supplemented with 20ng/ml FGF2 and 50ng/ml BMP4 changing medium every two days. For cardiac mesoderm (CM) formation, cells were cultured for 36 hours in CDM-BSA (without insulin) supplemented with 20ng/ml FGF2, 10µM LY294002, 10ng/ml BMP4 and 50ng/ml Activin A (Dr. Marko Hyvönen, Cambridge University). Subsequently cells were cultured for 4 days in CDM-BSA (without insulin) supplemented with 8ng/ml FGF2, 10ng/ml BMP4, 1µM IWR1 (WNT signalling inhibitor; Tocris Bioscience) and 0.5µM Retinoic Acid (Sigma-Aldrich), changing medium every two days. For presomitic mesoderm (PSM) formation, cells were cultured for 36 hours in CDM-BSA (without insulin) supplemented with 20ng/ml FGF2 and 8µM CHIR99021 (WNT signalling activator; Tocris Bioscience). Subsequently cells were cultured for 4 days in CDM-BSA (with insulin) supplemented with 4ng/ml FGF2, 1µM Retinoic Acid, 0.1µM LDN193189 (BMP signalling inhibitor; Sigma-Aldrich) and 10µM SB431542 (TGF-β signalling inhibitor; Tocris Bioscience). For mesoderm differentiation cells were plated on gelatin and mouse embryonic fibroblast (MEF) medium coated plates. Functional differentiation of the mesoderm subtypes was performed as described in the supplementary information.

### RNA extraction, cDNA synthesis and RT-qPCR

Total RNA was extracted using the GenElute™ Mammalian Total RNA Miniprep Kit (Sigma-Aldrich) and the On-Column DNase I Digestion set (Sigma-Aldrich) according to the manufacturer’s instructions. RNA was reverse transcribed using 250ng random primers (Promega), 0.5mM dNTPs (Promega), 20U RNAseOUT, 0.01M DTT and 25U of SuperScript II (all from Invitrogen). The resulting cDNA was diluted 30-fold for the qPCR reaction. Quantitative PCR mixtures were prepared using the KAPA SYBR^®^ FAST qPCR Master Mix (2X) Kit (Kapa Biosystems), 4.2µl of cDNA and 200nM of each of the forward and reverse primers. Samples were run on 384 well plates using the QuantStudio 12K Flex Real-Time PCR System machine and results analysed using the delta-delta cycle threshold method (ΔΔCt). Expression values were normalized to the housekeeping gene Porphobilinogen Deaminase (*PBGD*). Primer sequences are listed in supplementary table 1.

### Immunocytochemistry

Cells were fixed in 4% Paraformaldehyde (PFA) for 20 minutes at 4ºC followed by one wash with PBS. Cells were subsequently blocked and permeabilised at room temperature for 30 minutes in PBS with 4% donkey serum (Bio-Rad) and 0.1% Triton X-100 (Sigma-Aldrich). Primary antibodies were diluted in the same buffer and incubated with the cells overnight at 4ºC. After three washes with PBS, cells were incubated with AlexaFluor secondary antibodies for 1 hour at room temperature protected from light. Cells were subsequently washed three times for 5 minutes with PBS, adding Hoechst 33258 (bis-Benzimide H, 1: 10,000 dilution; Sigma-Aldrich) during the first wash to stain nuclei. The antibodies used are listed in supplementary table 2.

### Alcian blue staining

Monolayer cultures of chondrocytes were fixed in 4% paraformaldehyde (PFA) for 20 minutes at 4ºC. Cells were then washed with 0.5N HCl and stained overnight with 0.25% (w/v) Alcian Blue 8GX (Sigma-Aldrich) in 0.5N HCl. Stained cells were visualized using a Leica dissecting microscope. Alcian Blue dye was solubilized by overnight incubation with 8M guanidine hydrochloride (Sigma-Aldrich) and quantified by absorbance at 595nm using a spectrophotometer.

### Generation of CDK1 inducible knockdown line

Knockdown for CDK1 was performed using the single optimised inducible knockdown method as previously described (24, 53). Multiple shRNAs for the CDK1 gene were obtained from the validated shRNA database at Sigma-Aldrich. Briefly, shRNAs were introduced in the psOPTiKD plasmid between the BglII and SalI-HF sites. The psOPTiKD-shCDK1 vector was targeted to the AAVS1 locus by using 6μg of each of the following vectors: psOPTiKD-shCDK1, pZFN.AAVS1-KKR, and pZFN.AAVS1-ELD. H9 hESCs were nucleofected using the Lonza P3 Primary Cell 4D-Nucleofector X Kit and monoclonal colonies were selected for 7-10 days with 1 μg/ml of puromycin (Sigma-Aldrich). Tetracycline hydrochloride (Sigma-Aldrich) was used at 1μg/ml to induce the expression of the shRNA. Knockdown of CDK1 was confirmed by Western blot using anti-CDK1 antibody (Abcam, Ab133327). The shRNA sequences are listed in supplementary table 3. *Cellular fractionation and Western blot* - Cells were washed once with PBS and harvested with cell dissociation buffer (CDB; Gibco) for 10 minutes at 37^º^C. After one wash with cold 1% BSA-PBS, pellets were collected by centrifugation at 4ºC and 300g for 3 minutes. For isolation of the cytoplasmic fraction, pellets were resuspended in 6 times packed cell volume equivalent of Isotonic Lysis Buffer (10mM Tris-HCl, 3mM CaCl, 2mM MgCl_2,_ 0.32M sucrose, pH 7.5) supplement with protease and phosphatase inhibitors (Roche) and incubated for 12 minutes on ice. 0.3% Triton X100 (Sigma-Aldrich) was added for a further 3 minutes and samples were centrifuged at 1,800rpm for 5 minutes at 4ºC. The supernatant (cytoplasmic fraction) was collected in a fresh chilled tube. The nuclear pellet was washed once with Isotonic Lysis Buffer and centrifuged at 4,000rpm for 3 minutes at 4ºC. The nuclear pellet was resuspended in 2 times of the original packed cell volume equivalent of Nuclear Lysis Buffer (50mM Tris-HCl, pH 7.5, 100mM NaCl, 50mM KCl_2,_ 1mM EDTA, 10% Glycerol, 0.3% Triton X-100) supplemented with protease and phosphatase inhibitors (Roche), homogenized with a pellet pestle (Kimble Chase) and incubated for 30 minutes on ice. Subsequently, 125 units of benzonase nuclease (Sigma-Aldrich) was added to the lysates and incubated at room temperature for 45 minutes to remove nucleic acids. For extraction of whole cell lysate cells were lysed with RIPA lysis buffer (50mM Tris-HCl, 150mM NaCl, 1% Triton X-100, 0.5% sodium deoxycholate, 0.1% SDS, pH 8.0). Protein was quantified using the Pierce BCA Protein Assay Kit (Thermo Fisher Scientific) according to the manufacturer’s instructions. Lysates were prepared for Western blot by adding 1× NuPAGE LDS Sample Buffer (Thermo Fisher Scientific) with 1% β-mercaptoethanol and incubated at 95ºC for 5 minutes. Lysates (10-35 µg of protein) were electrophoresed on 12% NuPAGE Bis-Tris Precast Gels (Thermo Fisher Scientific) with NuPAGE MOPS SDS Running Buffer (Thermo Fisher Scientific). For identification of the size of the target protein, Precision Plus Protein Ladder was used (Bio-Rad). Protein was transferred on PVDF membranes (Bio-Rad) by liquid transfer using NuPAGE Transfer Buffer (Thermo Fisher Scientific). Membranes were blocked using 4% non-fat dried milk in Tris-buffered saline and 0.1% Tween (TBST buffer) for 30 minutes and probed with primary antibody overnight at 4ºC in TBST. Following 3 washes of 5 minutes each with TBST, membranes were incubated with horseradish peroxidase (HRP)-conjugated secondary antibody for 1 hour at room temperature in TBST. Membranes were then washed 3 times for 5 minutes with TBST and incubated with Pierce ECL Western Blotting Substrate and exposed to X-Ray Super RX Films (Fujifilm). In cases where Western blot membranes were incubated with more than one antibodies, the membranes were stripped using mild stripping buffer (1.5% Glycine, 0.1% SDS, 1% Tween 20, pH 2.2). Membranes were incubated twice for 10 minutes with the stripping buffer, twice for 10 minutes with PBS, twice for 5 minutes with TBST prior to blocking and incubation with primary antibody. The antibodies used are listed in supplementary table 4.

## Acknowledgements

This work was supported by the Wellcome Trust PhD program (PSAG/048 to L.Y.); the European Research Council advanced grant New-Chol (ERC: 741707 to L.V. and R.A.G), a BHF Senior Research Fellowship (FS/13/29/30024 to S.S.), a core support grant from the Wellcome and Medical Research Council to the Wellcome – Medical Research Council Cambridge Stem Cell Institute (PSAG028) and a core support grant from the Wellcome to the Wellcome Sanger Institute (WT206194).

## REFERENCES

1. Tam, P. P. L., and Loebel, D. A. F. (2007) Gene function in mouse embryogenesis: get set for gastrulation. Nat. Rev. Genet. 8, 368–381

2. Rodaway, A., and Patient, R. (2001) Mesendoderm: An ancient germ layer? Cell. 105, 169–172

3. Kimelman, D., and Griffin, K. J. (2000) Vertebrate mesendoderm induction and patterning. Curr. Opin. Genet. Dev. 10, 350–356

4. Arnold, S. J., and Robertson, E. J. (2009) Making a commitment: cell lineage allocation and axis patterning in the early mouse embryo. Nat. Rev. Mol. Cell Biol. 10, 91–103

5. Kinder, S. J., Tsang, T. E., Quinlan, G. A., Hadjantonakis, A., Nagy, A., and Tam, P. P. L. (1999) The orderly allocation of mesodermal cells to the extraembryonic structures and the anteroposterior axis during gastrulation of the mouse embryo. Development. 126, 4691–4701

6. Tam, P. P., Parameswaran, M., Kinder, S. J., and Weinberger, R. P. (1997) The allocation of epiblast cells to the embryonic heart and other mesodermal lineages: the role of ingression and tissue movement during gastrulation. Development. 124, 1631–42

7. Tam, P. P. L., and Trainor, P. A. (1994) Specification and segmentation of the paraxial mesoderm. Anat Embryol. 189, 275–305

8. Thomson, J. A., Itskovitz-Eldor, J., Shapiro, S. S., Waknitz, M. A., Swiergiel, J. J., Marshall, V. S., and Jones, J. M. (1998) Embryonic Stem Cell Lines Derived from Human Blastocysts. Science (80-.). 282, 1145–1148

9. Yiangou, L., Ross, A. D. B., Goh, K. J., and Vallier, L. (2018) Human Pluripotent Stem Cell-Derived Endoderm for Modeling Development and Clinical Applications. Cell Stem Cell. 22, 485–499

10. Theunissen, T. W., and Jaenisch, R. (2017) Mechanisms of gene regulation in human embryos and pluripotent stem cells. Development. 144, 4496–4509

11. Murry, C. E., and Keller, G. (2008) Differentiation of embryonic stem cells to clinically relevant populations: lessons from embryonic development. Cell. 132, 661–680

12. Gonzales, K. A. U., Liang, H., Lim, Y.-S., Chan, Y.-S., Yeo, J.-C., Tan, C.-P., Gao, B., Le, B., Tan, Z.-Y., Low, K.-Y., Liou, Y.-C., Bard, F., and Ng, H.-H. (2015) Deterministic Restriction on Pluripotent State Dissolution by Cell-Cycle Pathways. Cell. 162, 564–579

13. Pauklin, S., and Vallier, L. (2013) The cell-cycle state of stem cells determines cell fate propensity. Cell. 155, 135–47

14. Pauklin, S., Madrigal, P., Bertero, A., and Vallier, L. (2016) Initiation of stem cell differentiation involves cell cycle-dependent regulation of developmental genes by Cyclin D. Genes Dev. 30, 421–433

15. Singh, A. M., Sun, Y., Li, L., Zhang, W., Wu, T., Zhao, S., Qin, Z., and Dalton, S. (2015) Cell-Cycle Control of Bivalent Epigenetic Domains Regulates the Exit from Pluripotency. Stem Cell Reports. 5, 1–14

16. Singh, A. M., Chappell, J., Trost, R., Lin, L., Wang, T., Tang, J., Wu, H., Zhao, S., Jin, P., and Dalton, S. (2013) Cell-cycle control of developmentally regulated transcription factors accounts for heterogeneity in human pluripotent cells. Stem Cell Reports. 1, 532–544

17. Neganova, I., Zhang, X., Atkinson, S., and Lako, M. (2009) Expression and functional analysis of G1 to S regulatory components reveals an important role for CDK2 in cell cycle regulation in human embryonic stem cells. Oncogene. 28, 20–30

18. Neganova, I., Tilgner, K., Buskin, A., Paraskevopoulou, I., Atkinson, S. P., Peberdy, D., Passos, J. F., and Lako, M. (2014) CDK1 plays an important role in the maintenance of pluripotency and genomic stability in human pluripotent stem cells. Cell Death Dis. 5, e1508; doi:10.1038/cddis.2014.464

19. Bernardo, A. S., Faial, T., Gardner, L., Niakan, K. K., Ortmann, D., Senner, C. E., Callery, E. M., Trotter, M. W., Hemberger, M., Smith, J. C., Bardwell, L., Moffett, A., and Pedersen, R. A. (2011) BRACHYURY and CDX2 mediate BMP-induced differentiation of human and mouse pluripotent stem cells into embryonic and extraembryonic lineages. Cell Stem Cell. 9, 144–55

20. Mendjan, S., Mascetti, V. L., Ortmann, D., Ortiz, M., Karjosukarso, D. W., Ng, Y., Moreau, T., and Pedersen, R. A. (2014) NANOG and CDX2 pattern distinct subtypes of human mesoderm during exit from pluripotency. Cell Stem Cell. 15, 310–325

21. Cheung, C., Bernardo, A. S., Trotter, M. W. B., Pedersen, R. A., and Sinha, S. (2012) Generation of human vascular smooth muscle subtypes provides insight into embryological origin–dependent disease susceptibility. Nat. Biotechnol. 30, 165–173

22. Cheung, C., Bernardo, A. S., Pedersen, R. a, and Sinha, S. (2014) Directed differentiation of embryonic origin-specific vascular smooth muscle subtypes from human pluripotent stem cells. Nat. Protoc. 9, 929–38

23. Cheung, C., Bernardo, A. S., Trotter, M. W. B., Pedersen, R. A., and Sinha, S. (2012) Generation of human vascular smooth muscle subtypes provides insight into embryological origin-dependent disease susceptibility. Nat. Biotechnol. 30, 165–73

24. Bertero, A., Pawlowski, M., Ortmann, D., Snijders, K., Yiangou, L., Cardoso de Brito, M., Brown, S., Bernard, W. G., Cooper, J. D., Giacomelli, E., Gambardella, L., Hannan, N. R. F., Iyer, D., Sampaziotis, F., Serrano, F., Zonneveld, M. C. F., Sinha, S., Kotter, M., and Vallier, L. (2016) Optimized inducible shRNA and CRISPR/Cas9 platforms for *in vitro* studies of human development using hPSCs. Development. 143, 4405–4418

25. Harbour, J. W., and Dean, D. C. (2000) The Rb/E2F pathway: Expanding roles and emerging paradigms. Genes Dev. 14, 2393–2409

26. Yu, P., Pan, G., Yu, J., and Thomson, J. A. (2011) FGF2 Sustains NANOG and Switches the Outcome of BMP4-Induced Human Embryonic Stem Cell Differentiation. Cell Stem Cell. 8, 326–334

27. Vallier, L., Touboul, T., Chng, Z., Brimpari, M., Hannan, N., Millan, E., Smithers, L. E., Trotter, M., Rugg-Gunn, P., Weber, A., and Pedersen, R. a (2009) Early cell fate decisions of human embryonic stem cells and mouse epiblast stem cells are controlled by the same signalling pathways. PLoS One. 4, e6082

28. Olivera-Martinez, I., Harada, H., Halley, P. A., and Storey, K. G. (2012) Loss of FGF-Dependent Mesoderm Identity and Rise of Endogenous Retinoid Signalling Determine Cessation of Body Axis Elongation. PLoS Biol. 10.1371/journal.pbio.1001415

29. Ciruna, B., and Rossant, J. (2001) FGF Signaling Regulates Mesoderm Cell Fate Specification and Morphogenetic Movement at the Primitive Streak. Dev. Cell. 1, 37–49

30. Casey, E. S., O’Reilly, M. a, Conlon, F. L., and Smith, J. C. (1998) The T-box transcription factor Brachyury regulates expression of eFGF through binding to a non-palindromic response element. Development. 125, 3887–3894

31. Isaacs, H. V., Pownall, M. E., and Slack, J. M. W. (1995) eFGF is expressed in the dorsal midline of Xenopus laevis. Int. J. Dev. Biol. 39, 575–579

32. Isaacs, H. V, Pownall, M. E., and Slack, J. M. (1994) eFGF regulates Xbra expression during Xenopus gastrulation. EMBO J. 13, 4469–81

33. Griffin, K., Patient, R., and Holder, N. (1995) Analysis of FGF function in normal and no tail zebrafish embryos reveals separate mechanisms for formation of the trunk and the tail. Development. 121, 2983–94

34. Hu, Y., Feng, X., Mintz, A., Petty, W. J., and Hsu, W. (2016) Regulation of brachyury by fibroblast growth factor receptor 1 in lung cancer. Oncotarget. 7, 87124–87135

35. Teo, A. K. K., Arnold, S. J., Trotter, M. W. B., Brown, S., Ang, L. T., Chng, Z., Robertson, E. J., Dunn, N. R., and Vallier, L. (2011) Pluripotency factors regulate definitive endoderm specification through eomesodermin. Genes Dev. 25, 238–250

36. Radzisheuskaya, A., Le, G., Chia, B., Santos, R. L., Theunissen, T. W., Castro, L. F. C., Nichols, J., and Silva, J. C. R. (2013) A defined Oct4 level governs cell state transitions of pluripotency entry and differentiation into all embryonic lineages. Nat. Cell Biol. 15, 579–590

37. Loh, K. M., and Lim, B. (2011) A Precarious Balance: Pluripotency Factors as Lineage Specifiers. Cell Stem Cell. 8, 363–369

38. Brumbaugh, J., Russell, J. D., Yu, P., Westphall, M. S., Coon, J. J., and Thomson, J. A. (2014) NANOG is multiply phosphorylated and directly modified by ERK2 and CDK1 in vitro. Stem Cell Reports. 2, 18–25

39. Tsutsui, T., Hesabi, B., Moons, D. S., Pandolfi, P. P., Hansel, K. S., Koff, A., and Kiyokawa, H. (1999) Targeted disruption of CDK4 delays cell cycle entry with enhanced p27(Kip1) activity. Mol. Cell. Biol. 19, 7011–9

40. Rane, S. G., Dubus, P., Mettus, R. V, Galbreath, E. J., Boden, G., Reddy, E. P., and Barbacid, M. (1999) Loss of Cdk4 expression causes insulin-deficient diabetes and Cdk4 activation results in beta-islet cell hyperplasia. Nat. Genet. 22, 44–52

41. Berthet, C., Aleem, E., Coppola, V., Tessarollo, L., and Kaldis, P. (2003) Cdk2 Knockout Mice Are Viable. Curr. Biol. 13, 1775–1785

42. Malumbres, M., Sotillo, R., Santamaria, D., Galan, J., Cerezo, A., Ortega, S., Dubus, P., and Barbacid, M. (2004) Mammalian cells cycle without the D-type cyclin-dependent kinases Cdk4 and Cdk6. Cell. 118, 493–504

43. Santamaría, D., Barrière, C., Cerqueira, A., Hunt, S., Tardy, C., Newton, K., Cáceres, J. F., Dubus, P., Malumbres, M., and Barbacid, M. (2007) Cdk1 is sufficient to drive the mammalian cell cycle. Nature. 448, 811–815

44. Berthet, C., Klarmann, K. D., Hilton, M. B., Suh, H. C., Keller, J. R., Kiyokawa, H., and Kaldis, P. (2006) Combined Loss of Cdk2 and Cdk4 Results in Embryonic Lethality and Rb Hypophosphorylation. Dev. Cell. 10, 563–573

45. Diril, M. K., Koumar, C., Padmakumar, V. C., Du, T., and Wasser, M. (2012) Cyclin-dependent kinase 1 (Cdk1) is essential for cell division and suppression of DNA re-replication but not for liver regeneration. PNAS. 109, 3826–3831

46. Bienvenu, F., Jirawatnotai, S., Elias, J. E., Meyer, C. A., Mizeracka, K., Marson, A., Frampton, G. M., Cole, M. F., Odom, D. T., Odajima, J., Geng, Y., Zagozdzon, A., Jecrois, M., Young, R. A., Liu, X. S., Cepko, C. L., Gygi, S. P., and Sicinski, P. (2010) Transcriptional role of cyclin D1 in development revealed by a genetic–proteomic screen. Nature. 463, 374–378

47. Aleem, E., Kiyokawa, H., and Kaldis, P. (2005) Cdc2-cyclin E complexes regulate the G1/S phase transition. Nat. Cell Biol. 7, 831–6

48. Lee, T. I., Jenner, R. G., Boyer, L. A., Guenther, M. G., Levine, S. S., Kumar, R. M., Chevalier, B., Johnstone, S. E., Cole, M. F., Isono, K., Koseki, H., Fuchikami, T., Abe, K., Murray, H. L., Zucker, J. P., Yuan, B., Bell, G. W., Herbolsheimer, E., Hannett, N. M., Sun, K., Odom, D. T., Otte, A. P., Volkert, T. L., Bartel, D. P., Melton, D. A., Gifford, D. K., Jaenisch, R., and Young, R. A. (2006) Control of Developmental Regulators by Polycomb in Human Embryonic Stem Cells. Cell. 125, 301–313

49. Bracken, A. P., Dietrich, N., Pasini, D., Hansen, K. H., and Helin, K. (2006) Genome-wide mapping of Polycomb target genes unravels their roles in cell fate transitions. Genes Dev. 20, 1123–1136

50. Shen, X., Kim, W., Fujiwara, Y., Simon, M. D., Liu, Y., Mysliwiec, M. R., Yuan, G., Lee, Y., and Orkin, S. H. (2009) Jumonji Modulates Polycomb Activity and Self-Renewal versus Differentiation of Stem Cells. Cell. 139, 1303–1314

51. Wei, Y., Chen, Y.-H., Li, L.-Y., Lang, J., Yeh, S.-P., Shi, B., Yang, C.-C., Yang, J.-Y., Lin, C.-Y., Lai, C.-C., and Hung, M.-C. (2011) CDK1-dependent phosphorylation of EZH2 suppresses methylation of H3K27 and promotes osteogenic differentiation of human mesenchymal stem cells. Nat. Cell Biol. 13, 87–94

52. Yu, Y., Deng, P., Yu, B., Szymanski, J. M., Aghaloo, T., Hong, C., Wang, C.-Y., Tian, S., Hawkins, R. D., Leung, D., and al., et (2017) Inhibition of EZH2 Promotes Human Embryonic Stem Cell Differentiation into Mesoderm by Reducing H3K27me3. Stem Cell Reports. 8, 326–334

53. Bertero, A., Yiangou, L., Brown, S., Ortmann, D., Pawlowski, M., and Vallier, L. (2018) Conditional Manipulation of Gene Function in Human Cells with Optimized Inducible shRNA. Curr. Protoc. Stem Cell Biol. 44, 5C.4.1–5C.4.48

