## Supporting Information for "Cell cycle regulators control mesoderm specification in human pluripotent stem cells"

### Material Included

- Results: Figure S1
- Experimental procedures

### Results

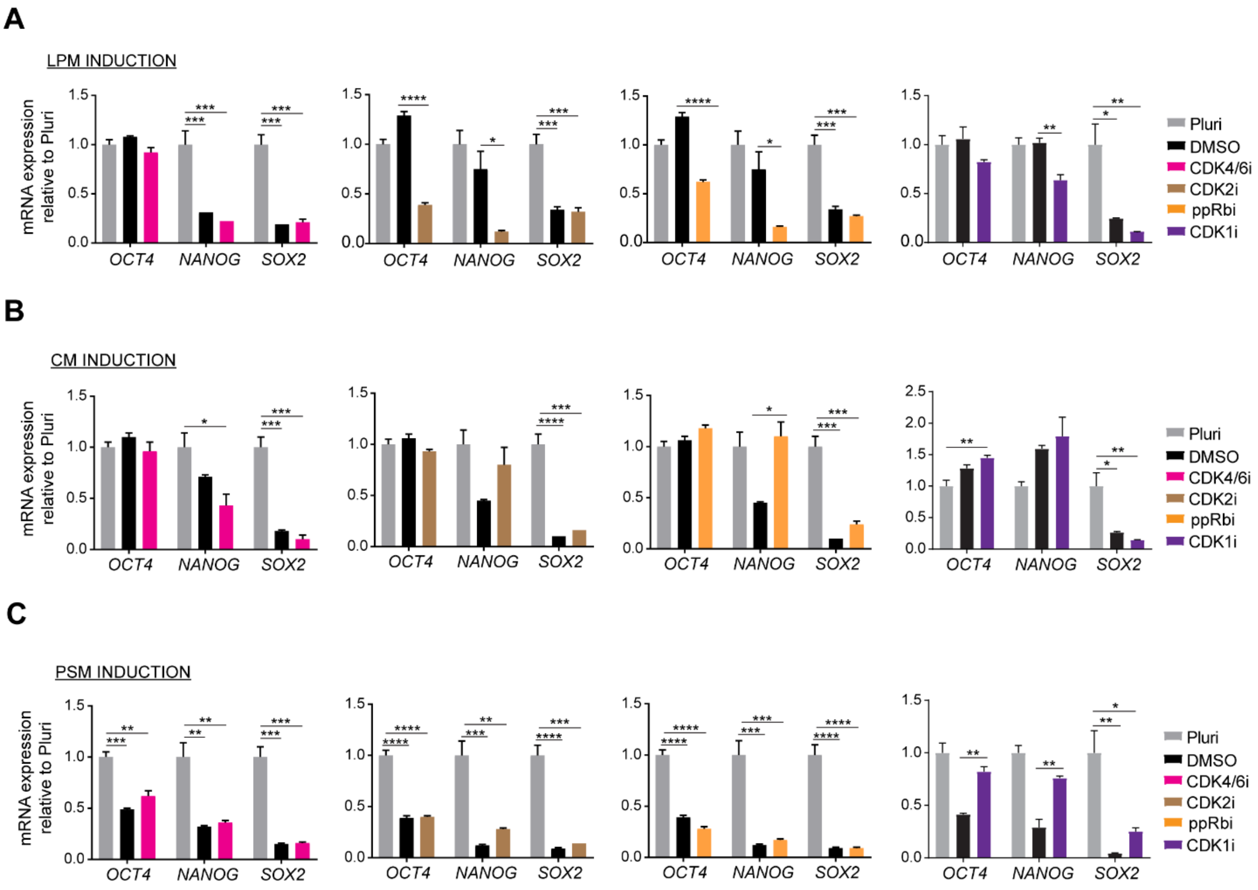


FIGURE S1. **Inhibition of G1 and G2/M cell cycle regulators affects pluripotency gene expression.** *A-C*, RT-qPCR analysis for expression of the pluripotency markers *OCT4*, *NANOG* and *SOX2* and *HAND1* during LPM induction (*A*), CM induction (*B*) and PSM induction (*C*) upon treatment with inhibitors of G1 and G2/M cell cycle regulators. Error bars represent ±SEM (n=3). Ordinary one-way ANOVA test followed by Tukey's test for multiple comparisons was performed. *p<0.05, **p< 0.01, ***p<0.001, ****p<0.0001.

### Experimental procedures

#### Table S1. qPCR primers used

| Gene | Forward Primer (5'-3') | Reverse Primer (5'-3') |
|---|---|---|
| *ACAN* | CCCCTGCTATTTCATCGACCC | GACACACGGCTCCACTTGAT |
| *ACTN1* | CAAACCTGACCGGGGAAAAAT | CTGAATAGCAAAGCGAAGGATGA |
| *CDX2* | GGCAGCCAAGTGAAAACCAG | TTCCTCTCCTTTGCTCTGCG |
| *CNN1* | GTCCACCCTCCTGGCTTT | AAACTTGTTGGTGCCCATCT |
| *COL2A1* | TGGACGCCATGAAGGTTTTCT | TGGGAGCCAGATTGTCATCTC |
| *EOMES* | ATCATTACGAAACAGGGCAGGC | CGGGGTTGGTATTTGTGTAAGG |
| *HAND1* | GTGCGTCCTTTAATCCTCTTC | GTGAGAGCAAGCGGAAAAG |
| *MIXL1* | GGTACCCCGACATCCACTTG | TAATCTCCGGCCTAGCCAAA |
| *NANOG* | CATGAGTGTGGATCCAGCTTG | CCTGAATAAGCAGATCCATGG |
| *NKX2.5* | GAGCCGAAAAGAAAGCCTGAA | CACCGACACGTCTCACTCAG |
| *PBGD* | GGAGCCATGTCTGGTAACGG | CCACGCGAATCACTCTCATCT |
| *T (BRACHYURY)* | TGCTTCCCTGAGACCCAGTT | GATCACTTCTTTCCTTTGCATCAAG |
| *TAGLN* | TCTTTGAAGGCAAAGACATGG | TTATGCTCCTGCGCTTTCTT |
| *TNNT2* | ACAGAGCGGAAAAGTGGGAAG | TCGTTGATCCTGTTTCGGAGA |

#### Table S2. Antibodies used for immunocytochemistry analysis

| Antibody | Species | Dilution | Catalogue Number | Manufacturer |
|---|---|---|---|---|
| BRACHYURY | Goat | 1:200 | AF2085 | R&D Systems |
| CARDIAC TROPONIN T | Rabbit | 1:500 | Ab45932 | Abcam |
| EOMES | Mouse | 1:200 | MAB6166 | R&D Systems |
| NKX2.5 | Goat | 1:500 | sc-14033 | Santa Cruz Biotechnology |
| PAX3 | Mouse | 1:100 | - | Dev. Hybridoma Bank |
| SM22a/TAGLN | Rabbit | 1:1,000 | ab-14106 | Abcam |
| Alexa Fluor 488 donkey anti-goat | - | 1:1,000 | A11055 | Invitrogen |
| Alexa Fluor 488 donkey anti-mouse | - | 1:1,000 | A21202 | Invitrogen |
| Alexa Fluor 488 donkey anti-rabbit | - | 1:1,000 | A21206 | Invitrogen |
| Alexa Fluor 647 donkey anti-goat | - | 1:1,000 | A21447 | Invitrogen |
| Alexa Fluor 647 donkey anti-mouse | - | 1:1,000 | A31571 | Invitrogen |
| Alexa Fluor 647 donkey anti-rabbit | - | 1:1,000 | A31573 | Invitrogen |

**Smooth Muscle Cell Differentiation**
For smooth muscle cell formation, LPM cells were dissociated with TrypLE Express (Life Technologies) for 5 minutes at 37ºC, washed once with CDM-PVA and centrifuged at 200g for 3 minutes. Cells were seeded on gelatin and MEF medium coated plates at a density of $2.6x10^4$ cells/$cm^2$ in CDM-PVA supplemented with 10ng/ml PDGF-BB (Peprotech) and 2ng/ml TGF-β (Peprotech) for 12 days. Media was changed every two days and cells split when confluent at a 1:2 ratio usually on day 3 or day 6.

**Cardiomyocyte Differentiation**
Following cardiac mesoderm formation, cells were cultured for two days in CDM-BSA (with insulin) supplemented with 8ng/ml FGF2 and 10ng/ml BMP4 (R&D) and subsequently fed every two days with CDM-BSA (with insulin). Onset of beating was observed on day 7-9 of differentiation.

**Chondrocyte Differentiation**
Following presomitic mesoderm formation, cells were cultured in CDM-BSA (with insulin) supplemented with 8ng/ml FGF2 and 10ng/ml BMP4 (R&D) for 10 days, changing media every two days.

Cells were treated with the small molecule inhibitors presented in table S3.

**Table S3. Small molecule cell cycle inhibitors used**

| **Inhibitor** | **Concentrations used** | **Catalogue Number** | **Supplier** |
|---|---|---|---|
| PD-0332991 | 5µM | S1116 | Selleckchem |
| RO-3306 | 10µM | S7747 | Selleckchem |
| Roscovitine | 4µM | R7772 | Sigma-Aldrich |
| RRD-251 | 10µM | R7532 | Sigma-Aldrich |

**Table S4. CDK1 shRNA sequences**

| **Top Oligo** | **Bottom Oligo** |
|---|---|
| GATCCCGCTGTACTTCGTCTTCTAATTCTCG AGAATTAGAAGACGAAGTACAGCTTTTTG | TCGACAAAAAGCTGTACTTCGTCTTCTAAT TCTCGAGAATTAGAAGACGAAGTACAGCGG |

**Table S5. Primary antibodies used for Western blot analysis**

| **Antibody** | **Species** | **Dilution** | **Catalogue Number** | **Manufacturer** |
|---|---|---|---|---|
| Phospho-Smad1/5 (Ser463/465) (41D10) | Rabbit (Biotinylated) | 1:500 | 9516S | Cell Signalling Technology |
| Smad1 | Rabbit | 1:500 | 9743S | Cell Signalling Technology |
| Phospho-β-catenin (Ser33/37/Thr41) | Rabbit | 1:1,000 | 9561S | Cell Signalling Technology |
| β-catenin | Goat | 1:1,000 | AF1329 | R&D Systems |
| Phospho-p44/p42 MAPK (Erk1/2) (Thr202/Tyr204) | Rabbit | 1:1,000 | 9101S | Cell Signalling Technology |
| p44/42 MAPK (Erk1/2) | Rabbit (Biotinylated) | 1:1,000 | 9102S | Cell Signalling Technology |
| A-tubulin | Mouse | 1:40,000 | T9026 | Sigma-Aldrich |
| B-actin | Mouse | 1:40,000 | A2228 | Sigma-Aldrich |
| Anti-CDK1 | Rabbit | 1:2,000 | Ab133327 | Abcam |
| Histone H3 | Rabbit | 1:40,000 | H0164 | Sigma-Aldrich |
| Anti-goat HRP | - | 1:10,000 | A5420 | Sigma-Aldrich |
| Anti-rabbit HRP | - | 1:10,000 | A0545 | Sigma-Aldrich |
| Anti-mouse HRP | - | 1:10,000 | A2554 | Sigma-Aldrich |
| Streptavidin-HRP | - | 1:2,000 | 3999S | Cell Signalling Technology |
